# Gene duplication drives genome expansion in Thaumarchaeota

**DOI:** 10.1101/2020.04.28.065540

**Authors:** Paul O. Sheridan, Sebastien Raguideau, Christopher Quince, Thames Consortium, Tom A. Williams, Cécile Gubry-Rangin

## Abstract

Ammonia-oxidising archaea of the phylum Thaumarchaeota are keystone species in global nitrogen cycling. However, only three of the six known families of the terrestrially ubiquitous order Nitrososphaerales possess representative genomes. Here we provide genomes for the three remaining families and examine the impact of gene duplication, loss and transfer events across the entire phylum. Much of the genomic divergence in this phylum is driven by gene duplication and loss, but we also detected early lateral gene transfer that introduced considerable proteome novelty. In particular, we identified two large gene transfer events into Nitrososphaerales. The fate of gene families originating on these branches was highly lineage-specific, being lost in some descendant lineages, but undergoing extensive duplication in others, suggesting niche-specific roles within soil and sediment environments. Overall, our results suggest that lateral gene transfer followed by gene duplication drives Nitrososphaerales evolution, highlighting a previously under-appreciated mechanism of genome expansion in archaea.

## Introduction

Thaumarchaeota have attracted much attention since their discovery, with accumulating evidence of their ubiquitous distribution and crucial ecosystem function in many environments^1-3^. Isolation and cultivation demonstrated that many of these organisms derive energy from the oxidation of ammonia under aerobic conditions. Ammonia oxidation is limited to a small number of microbial phyla but is one of the most important microbial metabolisms on the planet as it implements an essential and rate-limiting step in the nitrogen cycle, the conversion of ammonia to nitrite (via hydroxylamine). Within Thaumarchaeota, ammonia oxidising archaea (AOA) are ubiquitous and abundant in mesophilic soils and oceans, and are classically placed in four order-level phylogenetic lineages^1^: the Nitrososphaerales^4^, Nitrosopumilales^5^, *Candidatus* Nitrosotaleales^6^ and *Candidatus* Nitrosocaldales^7^.

While a substantial number of ecological studies and several cultures and enrichments confirm the ammonia-oxidising activity of Thaumarchaeota, there is also evidence that ammonia oxidation is not universal in this phylum^8-12^. In particular, there is evidence that the early-diverging lineages of Thaumarchaeota (including the Group 1.1c, Group 1.3 and pSL12 lineages) do not require ammonia oxidation for growth^9, 12^, and the ammonia oxidation machinery may have been acquired subsequent to the origin of Thaumarchaeota during, or after, the Great Oxygenation Event 2,300 My ago^11^. While oxygen and pH have been recognised as driving forces of massive diversification through their evolutionary history^11, 13^, the evolutionary mechanisms leading to such high phylogenetic and metabolic diversity have received little attention.

A large lateral gene transfer event in the last common ancestor (LCA) of AOA is proposed to have played a major role in their transition to a chemolithoautotrophic lifestyle^11, 14^. However, there is currently no evidence for other cases of large lateral gene acquisition in Thaumarchaeota evolution and little is known of the relative contributions of gene duplications and genes losses. Phylogenomic methods based on reconstructed ancestral gene contents have been used previously throughout the archaeal radiation to explicitly model gene family acquisitions, duplications and losses using a recently-developed approach for probabilistic gene mapping^15^ by amalgamated likelihood estimation (ALE)^16^. However, this study only included a limited number of thaumarchaeotal genomes. Similarly, a recent comparative analysis of a diverse set of thaumarchaeotal genomes^14^ did not include a large diversity of terrestrial thaumarchaeotal genomes nor disentangle the contributions of gene transfer and gene duplication to genomic diversification.

Here we present 12 high-quality metagenome-assembled genomes (MAGs), providing the first genome representatives of three of the six known families of the prevalent terrestrial order of AOA, the Nitrososphaerales. This enabled the reconstruction of a strongly supported phylogeny for the Nitrososphaerales, which was not previously achieved using single gene markers. Then, we investigated the mechanisms of genome evolution in the Thaumarchaeota phylum, quantifying the relative contributions of lateral gene transfers, gene duplications and gene losses. Phylogenomic analysis of the new genomes revealed two previously uncharacterised large lateral gene transfer events followed by subsequent extensive gene duplication mechanisms that have shaped the evolution of Nitrososphaerales.

## Results

### Thaumarchaeotal phylogenomic diversity

Phylogenomic analysis of 75 concatenated single-copy orthologues retrieved from 152 available thaumarchaeotal genomes (including 12 novel genomes from this study) (Table S1) provided a well-supported thaumarchaeotal phylogenomic tree (Figure 1), with most nodes with UF bootstrap values > 95 % and SH-aLRT values > 95 %. This phylogenomic tree was the best supported tree after comparison of several phylogenomic reconstruction methodologies (Supplementary Information: Extended phylogenomics). The tree is broadly similar to previously published work^11^ with some exceptions, mainly relating to poorly-supported basal branches in both trees (Supplementary Information: Extended phylogenomics).

**Figure 1.**
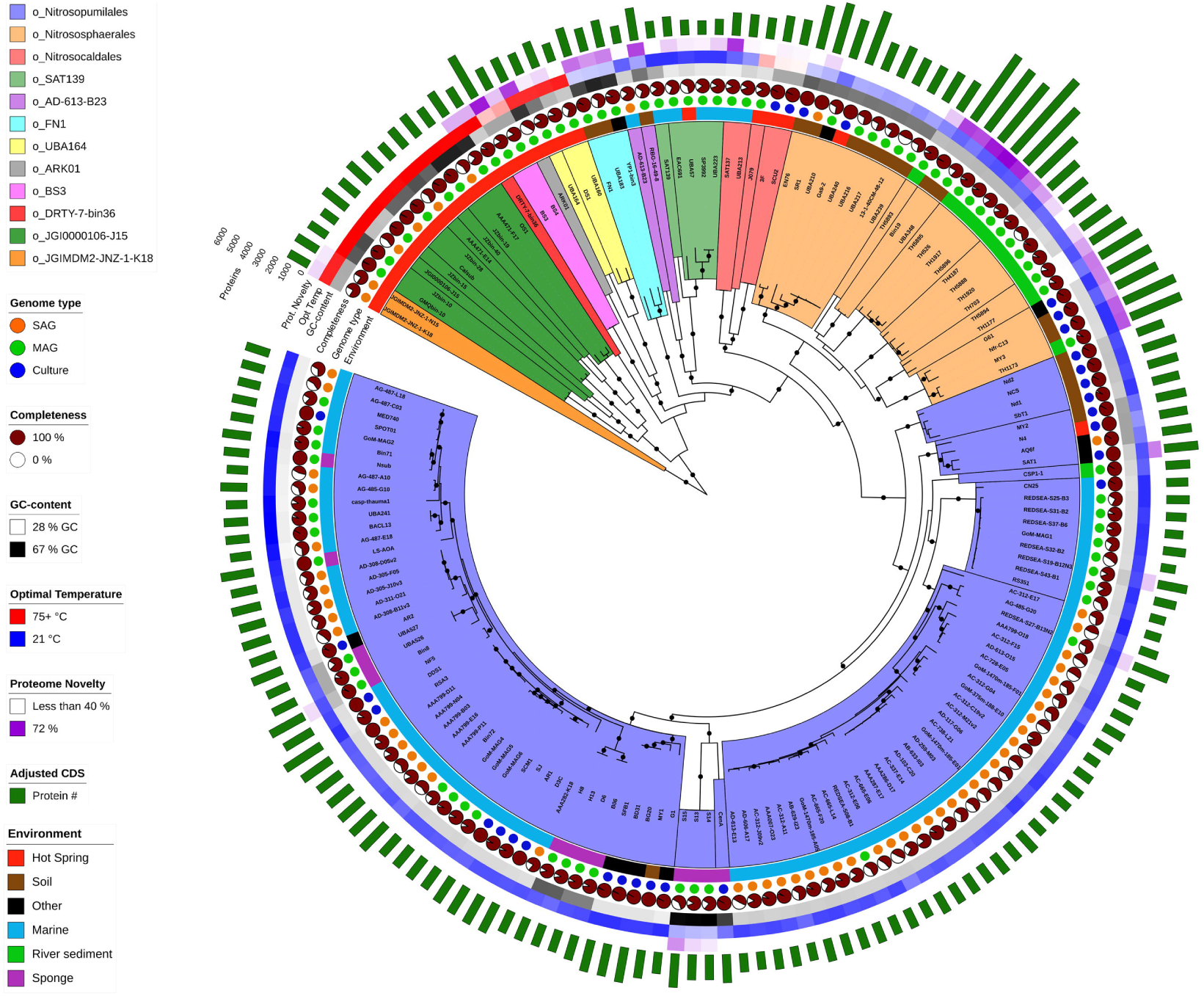
Phylogenomic tree of Thaumarchaeota. This tree includes 152 thaumarchaeotal genomes (29 culture genomes, 51 single-cell-assembled genomes (SAG) and 72 metagenome-assembled genome (MAGs), including twelve novel MAGs obtained in this study) from a range of environments and eleven Aigarchaeota genomes and is rooted with two Bathyarchaeota genomes. The tree was inferred by maximum likelihood from 75 concatenated phylogenetic markers, which were aligned separately and analysed using the best fitting model for each alignment. Dots indicate branches with greater than 95% ultrafast bootstrap and SH-aLRT support. Coloured order-level groups are further divided into families with black clade borders. CDS refers to coding sequences. Protein novelty is defined as the percentage of encoded proteins that lack a close homolog (e-value < 10-5, % ID > 35, alignment length > 80 and bit score > 100) in the arCOG database.

Previous classifications of Thaumarchaeota have been based on phylogenetic analyses of the ammonia monooxygenase (*amoA*) gene, but this gene is not present in all members of the phylum. We therefore suggest a phylum-wide thaumarchaeotal classification based on our phylogenomic analysis that maintains maximum consistency with previous work while incorporating the early-diverging lineages that lack *amoA*^2, 17, 18^ (Table S2). Our genome dataset of Thaumarchaeota represent a diverse phylum comprising 8 classes, 10 orders, 28 families, 31 genus and 103 species. While the classically used thaumarchaeotal nomenclature is congruent with the taxonomic stratification, a few exceptions were observed. For example, the order Nitrosopumilales encompasses *Candidatus* Nitrosotalea and Cenarchaeum, which were previously suggested to represent orders of their own, and Nitrososphaerales contains a minimum of 8 genera, with *Ca*. Nitrososphaera gargensis and *Nitrososphaera viennensis* and *Ca*. Nitrososphaera evergladensis belonging to different genera.

We also used the Genome Taxonomy Database Toolkit (GTDB-Tk) to evaluate the genomic diversity of our 12 new MAGs. While one of these genomes (TH1173) is a close relative of a published genome, MY3, the range of relative evolutionary divergence (RED)^19^ values for the other 11 MAGs was 0.72-0.82, suggesting that some of these MAGs represent new family-level lineages. These results are consistent with our phylogenomic analysis, in which these genomes represent novel deep branches within Nitrososphaerales (Figure 1). Previously, broad groups loosely reflecting the origins of thaumarchaeotal isolates and genomes were described as deep-water AOA, shallow-water AOA, terrestrial AOA and basal (non-ammonia oxidising) Thaumarchaeota^11^. Interestingly, assessment of their diversity at the genus level revealed that the deep-water AOA group is the least diverse, with 33 genomes affiliating to a single genus (max. divergence 26%), closely followed by the shallow-water AOA group, with 62 genomes forming 4 genera (max. divergence 44%). The terrestrial AOA represent the most diverse AOA group, with 40 genomes forming 15 genera (max. divergence 48%). However, the basal thaumarchaeotal group appears to be even more diverse with 17 genomes forming 11 genera (max. divergence 57%) (Figure S1).

The genomes used in this study presented relatively high completeness (especially metagenomes and culture genomes), with lowest values being reported for the single-cell genomes from the marine environment (Figure 1). Despite this methodological variation, and despite the wide range of genome size across the phylum (0.8 to 5.2 Mbp), there is a strong signal for higher genome size in some of the terrestrial Nitrososphaerales, in particular those within the genera g_Bin19 and g_TH5895. GC content is closely related to phylogeny in most cases, with a notable exception being the sponge-associated Thaumarchaeota, which present a higher GC content, potentially due to adaptation to environmental stresses such as nutrient and energy limitation^20^. Prediction of optimal growth temperatures (OGTs) confirmed that most of the Nitrososphaerales and Nitrosopumilales are mesophilic with OGTs between 21 and 37°C, with a few exceptions such as organisms affiliating to three closely related terrestrial genera (g_EN76, g_UBA210 and g_Nitrososphaera) or to two genera containing exclusively sponge-associated Thaumarchaeota (g_Cenarchaeum and g_S14) with slightly higher temperature optimum (range 33-42°C). As predicted, Nitrosocaldes have higher OGTs (range 32-56°C), but these estimates are lower than measured OGT in the two cultures available (difference of 15 and 19 °C, Supplementary Information: Extended optimal growth temperature). Predicted OGTs among the deeply-rooted non-AOA Thaumarchaeota are consistent with the hypothesis that Thaumarchaeota evolved from a hyperthermophilic common ancestor (Figure 1 and Table S1), but the presence of mesophiles in Nitrosocaldales and in their closest non-AOA relatives are consistent with the hypothesis that the last common ancestor of AOA was a mesophile, placing the hot origin of Thaumarchaeota further back in evolutionary history than previously proposed^21^.

### Phylogeny and metabolic traits of the Nitrososphaerales lineage

The 12 new MAGs presented in this work represent the first genome representatives of three of the six Nitrososphaerales families, enabling a detailed analysis of this order. As the whole-phylum analysis did not result in a highly supported phylogeny of the Nitrososphaerales order, a more targeted phylogenomic analysis based on 188 Nitrososphaerales markers was performed to increase resolution (Figure 2). With the addition these new MAGs, Nitrososphaerales consists of 22 species, representing 9 genera. The basal tree topologies based on the widely-used single genetic marker *amoA*^2, 13^ and on 188 single-copy core markers of this order are different, but a phylogenomically-informed rooting of the *amoA* phylogeny resolved many of the incongruencies (Figure S2). Additionally, the splitting of the families NS-alpha, NS-beta and NS-gamma, which could not be resolved using the *amoA* phylogeny alone, was resolved with high confidence (>95 % ultra-fast bootstrapping) using this phylogenomic approach. Moreover, the genome TH5893 appears to represent a previously undiscovered seventh family of Nitrososphaerales, but this remains unclear as its relationship to NS-gamma was not resolved with high confidence (Figure 2).

**Figure 2.**
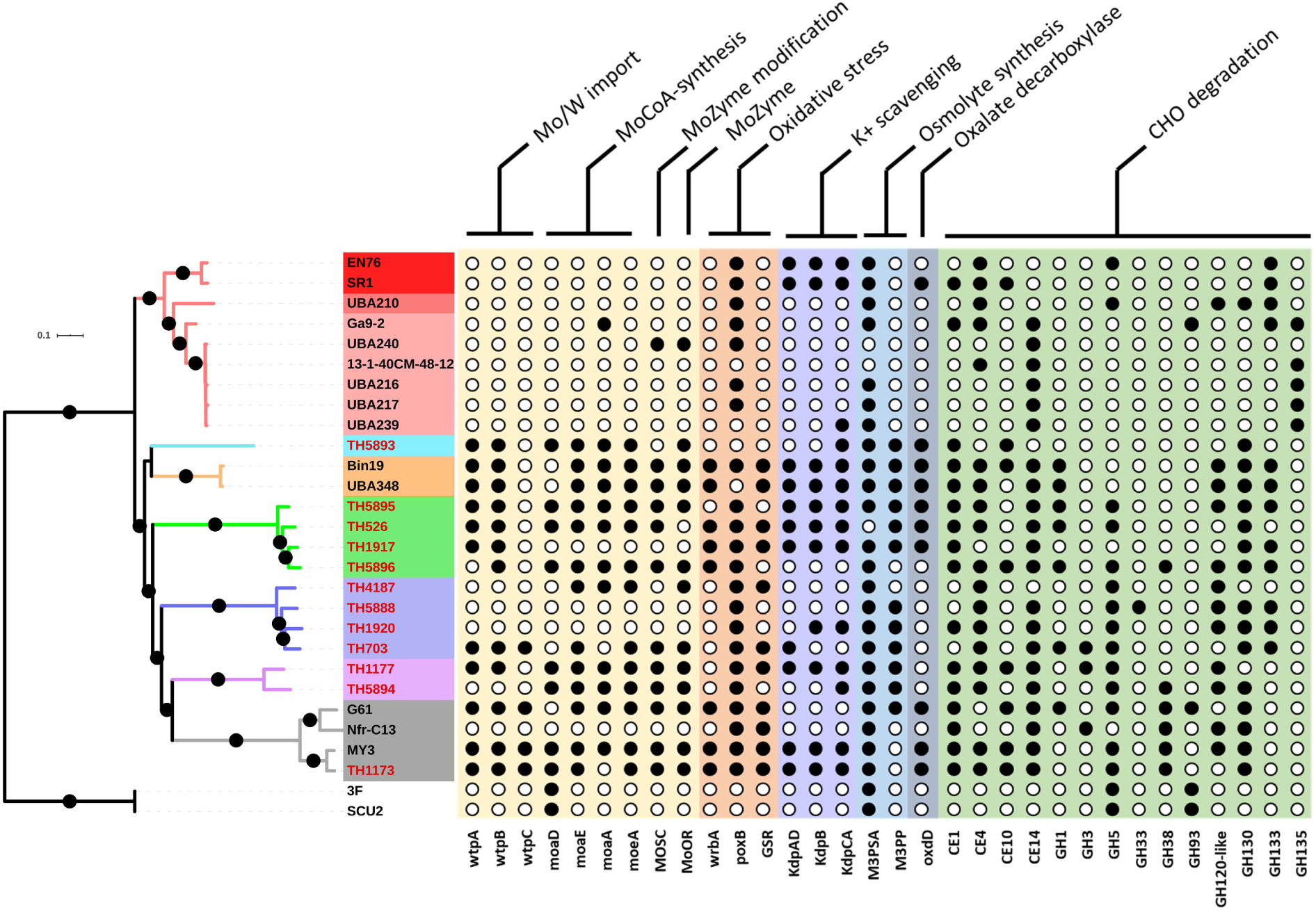
Phylogeny and distinctive metabolic traits of Nitrososphaerales. This tree includes 26 Nitrososphaerales genomes and was inferred by maximum likelihood from 188 concatenated marker genes. The twelve new genomes obtained in this work are indicated by red tip labels and the tree is rooted with the *Ca*. Nitrosocaldales strains 3F and SCU2. This tree has similar topology to the phylum-wide tree, but with strong support for almost all branches. Dots indicate branches with greater than 95% ultrafast bootstrap and SH-aLRT support. Colours on branches and leaf labels indicate families and genera, respectively. While many of the genes acquired in early stages of Nitrososphaerales evolution are uncharacterised, several genes involved in molybdoenzymes synthesis and stress response could be identified. The following genes were used in the figure: *wtpABC*, Wtp transport system subunits A, B and C; MOSC, molybdenum cofactor sulfurase; MoOR, Molybdopterin oxidoreductase; *wrbA*, NAD(P)H quinone oxidoreductase; *poxB*, pyruvate oxidase; GSR, glutathione-disulfide reductase; *kdpABC*, potassium transport system subunits A, B and C; M3PSA, mannosyl-3-phosphoglycerate synthase; M3PP, mannosyl-3-phosphoglycerate phosphatase; CE, carbohydrate esterase family; GH, glycoside hydrolase family.

AOA are a highly metabolically conserved group of chemolithoautotrophic organisms, which use ammonia for energy22 and fix CO_2_ as a carbon source23. They also have the capacity to synthesise many of the cofactors^24^ and amino acids required for their cellular function. Despite this high level of metabolic conservation, Nitrososphaerales AOA possess additional metabolic capacities that likely facilitated their expansion into a wide variety of soil, wastewater and river sediment environments^25^ (Figure 2). Among the traits acquired early in the Nitrososphaerales diversification, many likely contribute to protection against osmotic and oxidative stress, production of molybdoenzymes and carbohydrate utilisation.

Carbohydrate active enzymes (CAZymes) are ubiquitous in the tree of life, allowing organisms to degrade environmental carbohydrates as energy and carbon sources (such as the utilisation of monosaccharides from cellulose or hemicellulose degradation) and for the formation of more complex compounds from simpler substrates (such as extracellular polysaccharide synthesis)^26^. The number of CAZyme gene acquisitions in the Nitrososphaerales lineage is large and greater than that of other members of the phylum, as the other Thaumarchaeota encode an average of less than two CAZymes. The last common ancestor of the Nitrososphaerales is predicted to have acquired the amylo-α-1,6-glucosidase GH133 (EC3.2.1.33) and polyspecific carbohydrate esterases CE1, CE4 and CE10. Additionally, the beta-mannanase GH130, alpha-mannanase GH38, hemicellulases GH1, GH3 and GH5, and a putative chitin disaccharide deacetylase (CE14) were also acquired during Nitrososphaerales diversification.

Since these organisms lack the phosphofructokinase necessary to complete the glycolytic pathway, it is difficult to determine the purpose of these carbohydrate degrading enzymes. One possible explanation is that the resulting monosaccharides are being utilised for the biosynthesis of cellular components such as extracellular polysaccharides or osmolytes. Indeed, the Nitrososphaerales possess mannosyl-3-phosphoglycerate synthase and mannosyl-3-phosphoglycerate phosphatase genes, giving these organisms the potential ability to synthesise the osmoprotectant mannosyl-glycerate from mannose monosaccharides and glycerate 3-phosphate. They have also lost the mannose-6-phosphate isomerase that is present in most other Thaumarchaeota, preventing mannose monosaccharides being shunted into early glycolysis, increasing the amount of mannose available for mannosyl-glycerate synthesis.

Kdp, a high affinity ATP-driven potassium uptake system that enables K^+^-mediated osmoregulation in potassium limited environments^27^, is present in several Nitrososphaerales genomes, but almost completely absent from marine Thaumarchaeota. In the event of osmotic shock, this system could allow the organisms to maintain turgor pressure in soils where there is low potassium concentration or where potassium is bound to negatively charged humus or clay particles by cation exchange.

Nitrososphaerales have also acquired several systems for protection against oxidative stress indicating that there may be additional production of reactive oxygen species (ROS) in the metabolism of this lineage. These include pyruvate oxidase (*PoxB*), NAD(P)H quinone oxidoreductase (*WrbA*) and glutathione-disulphide reductase (*GSR*). *PoxB* catalyses the ubiquinone-dependent oxidative decarboxylation of pyruvate to form acetate and CO_2_. This pathway enables the production of acetate in a NAD-independent manner, preventing the accumulation of ROS from resulting NAD+ regeneration by NADH-dehydrogenase in slow growing cells where ROS are not diluted by cell growth^28^. *WrbA* maintains quinones in the fully reduced state, guarding against the production of ROS from one-electron redox cycling ^29^ and *GSR* catalyses the reduction of glutathione disulphide to two reduced molecules of glutathione – an antioxidant that protects cellular components from oxidative stress.

Another notable difference between the Nitrososphaerales and other lineages of AOA is the presence of genes for the uptake of molybdate and synthesis of molybdoenzymes (MoZymes). MoZymes are present in a wide range of eukaryotes and prokaryotes in many diverse environments where they play crucial roles in detoxification, nitrate assimilation and anaerobic respiration as redox reaction catalysts that transfer electrons to a range of terminal electron acceptors^30^. While MoZymes have been reported in basal members of the Thaumarchaeota, this is the first report of these enzymes in AOA.

The *Wtp* transport system, which has high affinity for both molybdate and tungstate, is present in many Nitrososphaerales, but almost completely absent in other Thaumarchaeota. The enzymes *MoaA, MoaD/E* and *MoeA* catalyse the formation of molybdenum cofactor (MoCo) of MoZymes and MOSC transfers sulfur to the molybdenum, forming mono-oxo MoCo^31^. MoZymes of the sulphite oxidase (SO) superfamily were identified in several members of the Nitrososphaerales order (Figure S3). These enzymes appear to be an intermediate between the eukaryotic SO and the bacterial protein-methionine-sulfoxide reductase (Figure S3). The majority of the Nitrososphaerales MoZymes are structurally similar to the eukaryotic SO, possessing both the oxidoreductase molybdopterin binding domain (PF00174) and the MoCo oxidoreductase dimerisation domain (SSF81296), but lacking the N-terminal cytochromes b5 electron transport hemoprotein domain (PF00173). These enzymes however possess at their N-terminal several transmembrane helicases that are likely to be a novel class of cytochrome domains. This protein family is therefore likely to be a class of archaeal SO that convert sulphite to sulphate, allowing the generation of ATP in oxidative phosphorylation. This reaction produces hydrogen peroxide as a by-product, possibly explaining the need for extra oxidative stress protection mechanisms in this order.

The three MoZymes of s_Bin19 and one of the two MoZymes of *Ca*. Nitrosocosmicus oleophilus (MY3_01316) were more similar in structure and sequence to the bacterial protein-methionine-sulfoxide reductase that repairs oxidized periplasmic proteins containing methionine sulfoxide residues, indicating a putative role of this enzyme in oxidative stress damage protection.

### Influence of evolutionary mechanisms on thaumarchaeotal diversification

The exhaustive creation of probabilistic ancestral reconstructions from every branch of the thaumarchaeotal phylogeny allowed the characterisation and quantification of proteome changes along every lineage (Figure 3 and S5; Table S4). The majority of the predicted 67,400 gene content gains in Thaumarchaeota evolution occurred through 51,653 duplications (77% of gains) of pre-existing genes, with 11,430 intra-phylum gene transfers (intra-LGT) (17% of gains) and 4,317 originations (including inter-phylum gene transfers (inter-LGT) and *de novo* gene formation) (6% of gains). There were also 227,837 gene losses predicted, indicating gene duplication and gene loss as the two most significant drivers of gene content change in Thaumarchaeota evolution. While there were nearly 3.5-fold more losses than gains, most of the losses occurred on the recent branches, explaining their preponderance.

**Figure 3.**
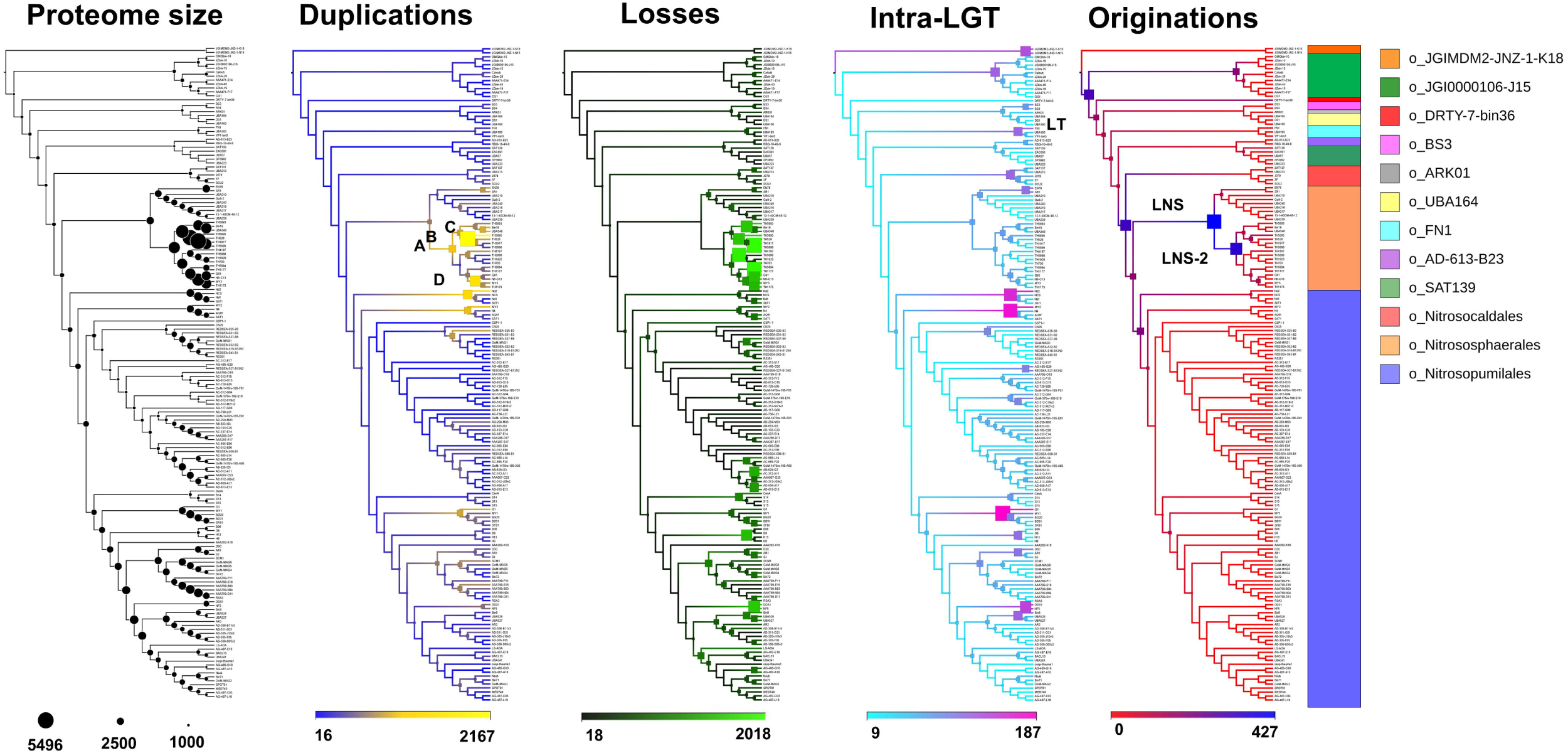
Quantified mechanisms of proteome evolution. Proteome changes were predicted on each branch of the thaumarchaeotal phylogeny. Duplication and loss are defined as the copying and loss of a gene within a genome, respectively. Intra-LGT is defined as the acquisition of a gene from other member(s) of the phylum, while originations are defined as the acquisition of a gene from members of other phyla outside the sampled genome set (Inter-LGT) and possibly by *de novo* gene formation. Scale numbers indicate the range of the predicted number of events for a given mechanism. The cladogram possesses the topology of the ML tree presented in Figure 1. Coloured bar indicates order-level classification of tree leaves. Originations occurring in the last common ancestors of all Thaumarchaeota, Nitrososphaerales and a multifamily sub-group within Nitrososphaerales are labelled LT, LNS and LNS-2 respectively. A, B, C and D indicate branches within the Nitrososphaerales order with extensive duplication.

Four branches within the Nitrososphaerales order (noted A, B, C and D in Figure 3) are estimated to undergo particularly high rates of gene duplication, collectively accounting for 11% of duplication events in the phylum (Figures 3, S5 and Table S4), with more than 900 duplication events occurring on each of those branches. Numerically, these are the major mechanisms of predicted proteome expansion in the Thaumarchaeota phylum, resulting in most of Nitrososphaerales encoding over 3,000 proteins and possessing an average genome size of 2.7 Mbp (range 1.5 – 5.2 Mbp). Duplication of existing genes also appears to be the major mechanism of predicted proteome expansion in the *Ca*. Nitrosotalea and *Ca*. Nitrosotenuis genera with their last common ancestors undergoing 1,337 and 1,010 duplication events, respectively (Table S4). This resulted in the extant members of these genera possessing the largest proteomes in the Nitrosopumilales order.

These duplication hotspots resulted in the expansion of several arCOG families (Table S5), with copy numbers increasing more than 10-fold in several families predicted to be involved in transcription (arCOG04362, arCOG01760 and arCOG01055), signal transduction (arCOG02391), carbohydrate transport (arCOG00144) and coenzyme metabolism (arCOG00972) (Table S5). These expansions also coincided with contractions of several other arCOGs (Table S6). Interestingly, the three most notable contractions in NS-beta LCA (arCOG01067, arCOG04946 and arCOG00041) (Table S6) were expanded in its ancestor, Branch 280 (Table S5), indicating a fluctuating expansion and contraction of families throughout evolutionary history, rather than a consistent selective direction.

The evolution of the Nitrososphaerales lineage appears to have involved a significantly larger number of proteome changes than the phylum as a whole (*P* < 1e-6), with the greater numbers of duplications (*P* < 0.006) and losses (*P* < 5e-5) explaining much of this difference.

Although duplications are the major mechanism of proteome expansion in Thaumarchaeota, acquisition of a large number of gene families from outside the phylum have also largely contributed. This is particularly true for the LCA of ammonia-oxidising Thaumarchaeota (LA), where the origination of 320 genes families contributed to the transition to a chemolithoautotrophic ammonia-oxidising lifestyle (Figure 3 and Table S4). Indeed, large gene family gains in LA have also been predicted in earlier studies^11, 14^. All major origination events (182 to 427 gene families) have occurred early in the evolutionary history of ammonia oxidising Thaumarchaeota and accounted for 48% of all gene family originations in the Nitrososphaeria. In addition, large gene family origination (166 gene families) also occurred in the last common ancestor of the entire thaumarchaeotal phylum (LT in Figure 3).

Two large origination events were predicted in the Nitrososphaerales introducing 427 and 353 new gene families into the order (on branches 305 and 280, respectively, Figure 4), making these the two largest origination events detected in the phylum (Figure 3). The majority of the genes originating on these branches (71 % and 83 % on branches 305 and 280, respectively, Table S7) lack close matches in the arCOG database^11^, reflecting a high level of proteome novelty (Figure 1). The source of these originating gene families may be inter-LGT or *de novo* gene formation. Homologs of the majority of these gene families (84 % and 59 % on branches 305 and 280, respectively, Table S8) were detected in a larger database of archaea, bacteria and eukaryote genomes (described in Materials and Methods), indicating their lateral acquisition. Most gene families acquired by the last common ancestor of Nitrososphaerales (LNS in Figure 3) are predicted to have been transferred laterally from other archaea (archaea 95%, bacteria 5%) (Table S8), whereas gene families acquired in the second large Nitrososphaerales LGT event (LNS-2 in Figure 3) were probably transferred from a variety of archaea (64%), bacteria (34%) and eukaryotes (2%) (Table S8). Gene transfer from other archaea was also demonstrated in the sister phylum Aigarchaeota^14^.

**Figure 4.**
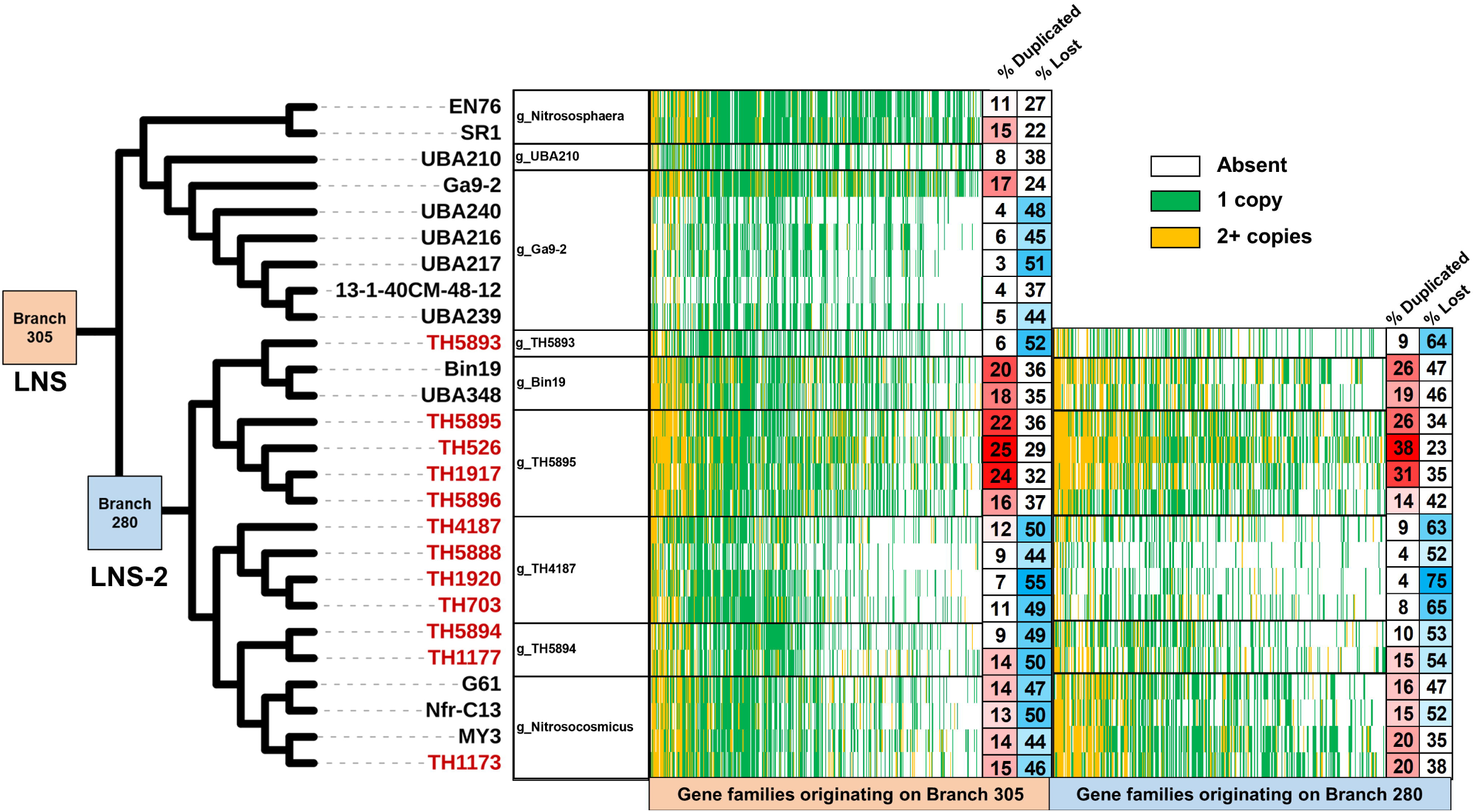
Fates of the originating gene families in Nitrososphaerales lineages: evidence for subsequent extensive duplication and loss. The cladogram highlights branches that experienced large origination events, LNS and LNS-2. The originating gene families are extensively duplicated in some lineages, such as g_TH5895, but also extensively lost in other lineages, such as g_TH4187. Colours indicate the fate of the originating gene families: duplicated (yellow), lost (white) or retained as a single copy (green). The ‘% Duplicated’ and ‘% Lost’ columns at the ends of the two heatmaps indicate the percentage (corrected for genome completeness) of originating gene families that underwent duplication or loss in a given extant member of the order. The 12 new genomes assembled in this study are indicated by red tip labels. Genus-level taxonomic affiliations are presented next to cladogram tips.

These originating gene families encountered different fates in the different lineages of Nitrososphaerales (Figure 4), with some families being subject to duplications or losses. Interestingly, 47 and 54% of originating gene families on these two branches were duplicated within at least one extant member of this order (Figure 4). The combination of these gene family originations followed by intensive duplications have resulted in members of the Nitrososphaerales having among the highest levels of proteome novelty in the Thaumarchaeota (Figure 1). A large number of losses (up to 75% of the originating gene families) occurred in g_TH4187 and g_Ga9-2 (Figure 4). Conversely, there was intensive duplication of originating gene families (up to 29% of the originating gene families) in the g_TH5895 and g_Bin19, the genera with the largest genomes in the phylum, indicating that originating genes are duplicated rather than lost in cases of proteome expansion. Similar trends of extensive loss in g_TH4187 and g_Ga9-2, and duplication in g_TH5895 and g_Bin19 were observed in ancestral gene families (originating prior to the LCA of Nitrososphaerales) (Table S9), but were significantly higher for originating gene families in g_TH4187 (*P* < 0.01), g_Ga9-2 (*P* < 0.01) and g_TH5895 (*P* < 0.05). The lineage-specific nature of gene duplication and loss of these originating gene families also suggests that they may be more important for specialisation of the different Nitrososphaerales lineages into distinct ecological niches than for more general soil and sediment environmental colonisation.

### Major transitions in thaumarcheotal evolution

Thaumarchaeota proteomes have undergone extensive gene content changes during their evolution, with some important niche transitions occurring. The nature and influence of these changes were analysed through ancestral reconstruction of the metabolic pathways at key nodes in the thaumarchaeotal phylogeny (Table S10).

The most extreme transition in the history of Thaumarchaeota was probably the transition from the LCA of Thaumarchaeota (LT) to the LCA of ammonia oxidising Thaumarchaeota (LA). Based on the genomic predictions, LT was a heterotrophic hyperthermophilic organism utilising carbohydrates as energy and carbon sources, with a complete glycolytic pathway (Table S10). The absence of genes for biosynthesis of cobalamin, riboflavin and biotin indicate that it acquired vitamins from extracellular organic sources. The transition from the LT to LA was accompanied by several functional gain and loss events (Table S11 and S12). Most notably, this transition involved the acquisition of the ammonia monooxygenase genes (*amoABC*), which generate energy by oxidising ammonia to hydroxylamine. This event coincided with the acquisition of urease (EC:3.5.1.5), which converts urea to ammonia, and genes for the biosynthesis of cobalamin, riboflavin and biotin. LA also acquired the ability to biosynthesis asparagine, alanine and threonine from aspartate through the acquisition of aspartate 4-decarboxylase (EC:4.1.1.12), asparagine synthase (EC:6.3.5.4), aspartate kinase (EC:2.7.2.4) and threonine synthase (EC:4.2.3.1) (Table S11 and S12).

The evolutionary transition from LT to the ammonia oxidising lifestyle of LA involved the loss of glucokinase (EC:2.7.1.2) and phosphofructokinase (EC:2.7.1.11), preventing them from generating ATP by glycolysis. They also lost the glycolysis-associated carbohydrate degrading enzymes alpha-N-arabinofuranosidase (EC:3.2.1.55), alpha-amylase (EC:3.2.1.1), alpha-mannosidase (EC:3.2.1.24) and the CO_2_ fixing genes phosphoenolpyruvate carboxylase (EC:4.1.1.31) and pyruvate ferredoxin oxidoreductase (EC:1.2.7.1). These losses were concomitant with the acquisition of the 3-hydropropionate/4-hydroxybutrate CO_2_ fixation pathway genes methylmalonyl-CoA epimerase (EC:5.1.99.1) and 4-hydroxybutyryl-CoA dehydratase (EC:4.2.1.120) (Table S11 and S12)^23^.

The establishment of LA as a chemolithoautotroph was followed by diversification into 3 order-level groups: Nitrosocaldales, present in hot spring and marine environment; Nitrososphaerales, present mostly in river sediments and soil; and Nitrosopumilales present mostly in marine, soil and sponge-associated environments.

The transition from LA to the last common ancestor of Nitrosocaldales (LNC) involved several functional gains, including geranylgeranylglycerol-phosphate geranylgeranyltransferase (EC:2.5.1.42), which is involved in the formation of polar membrane lipids in many thermophilic archaea, and the vitamin-B12-independent methionine synthase (EC:2.1.1.14), functionally replacing the vitamin-B12-dependent methionine synthase (EC:2.1.1.13), which is absent from every member of this order. The DNA polymerase D (EC:2.7.7.7) has been lost in LNC. Both the presence of vitamin-B12-independent methionine synthase and the absence of DNA polymerase D were previously reported in *Nitrosocaldus islandicus* 3F^32^, but present results expand those findings to all members of the order. Other notable losses in this transition are the Kdp, high affinity ATP-driven K+-transport system (EC:3.6.3.12), which is involved in osmotic stress resistance in low potassium environments and Uvr excinuclease, which is involved in DNA repair from ultraviolet DNA damage. Kpd is absent from almost all marine and hot spring Thaumarchaeota in this analysis, indicating that this system is not essential in aqueous environments. It has been proposed that the absence of Uvr in deep-water Nitrosopumilales is due to the lack of light in this environment^11^. This theory agrees with the absence of these genes in Nitrosocaldales, which derive from hot spring (in the case of 3F, SCU2 and J079) and deep ocean (in the case of SAT137 and UBA231) environments in which light is also likely to be absent.

During the transition from LA to the last common ancestor of Nitrososphaerales (LNS), many genes were acquired including the mismatch repair genes DNA adenine methylase (EC:2.1.1.72) and DNA helicase II (EC:3.6.4.12), possibly conferring additional resistance to DNA damage as these organisms expanded into environments with more light exposure such as surface soil and sediment. This transition also involved the loss of the *Phn* phosphonate transport system, which provides a source of phosphate in environments where alternative phosphorus sources are scarce^33^, which may be required less in more eutrophic soil and sediments (Table S11 and S12).

The evolution of LA to the last common ancestor of Nitrosopumilales (LNP) involved gain of ubiquinone-dependent (EC:7.1.1.2) and ubiquinone-independent (EC:1.6.99.3) NADH dehydrogenases (Table S11 and S12), giving LNP all three families of respiratory NADH dehydrogenases, which regenerate NAD+ from NADH by catalysing the transfer of electrons from NADH to coenzyme Q10 in oxidative phosphorylation. The advantage of encoding all three families remains largely unknown^34-36^, but could provide a conservation of respiratory function under changing metabolic conditions.

## Discussion

As for many prokaryotic lineages, the metabolic diversity of Thaumarchaeota remains to be fully characterized, especially in underexplored environments. The new genomes presented in this work offer the first genome representatives for three of the six known families of Nitrososphaerales. This enabled a resolution of Nitrososphaerales phylogeny that was not possible from *amoA* gene-based phylogenies and provided insights into their predicted physiology. Although most of the presently available thaumarchaeotal genomes belong to the marine groups, the majority of thaumarchaeotal phylogenetic diversity is present in non-ammonia oxidising Thaumarchaeota and in AOA from terrestrial environments. In addition to phylogenetic diversity, substantial genome novelty is also predicted in these organisms, with the Nitrososphaerales possessing the most novel gene families in the ammonia oxidising Thaumarchaeota, resulting from two large lateral gene transfer events in their history. The transition of Thaumarchaeota to an autotrophic ammonia-oxidising lifestyle has been proposed to result from a large lateral gene transfer event in the last common ancestor of AOA^11^. The new genomes presented in this study enabled the detailed analysis of the evolutionary history of Nitrososphaerales gene families, identifying two additional LGT events associated with large gene gains and extensive molecular innovation.

It has been proposed that archaeal genome evolution is driven by punctuated episodes of extensive gene acquisition, followed by lineage-specific gene loss^37^. While our analysis indicates that gene transfer also contributed to the evolution of Nitrososphaerales, the predominant mode of genome expansion was gene duplication of both ancestral and originating gene families, with higher duplication rates occurring in the latter. In light of this model, reassessment of a previously archaea-wide evolutionary analysis^15^ indicates that a similar pattern of large origination events and subsequent duplications also occurred in other archaeal lineages, especially for the lineages with the largest known archaeal genomes (Lokiarchaeum, *Haloarcula marismortui* and *Haloferax volcanii*). Therefore, our analysis of thaumarchaeotal evolution may have uncovered a previously unappreciated paradigm of genome expansion with broader significance. Larger genome sizes have been reported for mesophilic soil prokaryotes compared to marine relatives^38^, but it is unclear whether this is due to genome expansion in soil lineages from ancestors with smaller genomes, or from genome reduction in marine lineages. This work provides an example of a case in which soil-marine genome size differences are driven by genome expansion in soil lineages.

## Materials and Methods

### Sequence database

Thaumarchaeotal genomes were searched in literature and downloaded from NCBI (www.ncbi.nlm.nih.gov) and IMG (https://img.jgi.doe.gov/) genome reference databases. In addition, twelve novel MAGs were assembled based on a massive co-assembly approach performed on 171 sediments from the Thames river (see Supplementary Information). Thirteen diverse genomes of the most closely related archaea (eleven Aigarchaeota and two Bathyarchaeota) were also added to this database to allow rooting of the phylogenetic tree. All genomes were annotated using Prokka v1.11^39^ to ensure uniformity between annotation methods and genome completeness and contamination were estimated using CheckM^40^. Only genomes with completeness higher than 45% and presenting less than 10% contamination were kept in the final dataset (to avoid bias in phylogenomic analyses), reaching a total of 152 thaumarchaeotal genomes (29 culture genomes, 51 single-cell genomes and 72 MAGs (Table S1)). Several genome characteristics were estimated, including GC content (using QUAST^41^), total predicted genomic size (measured genome size corrected by the completeness score) and predicted optimal growth temperature (based on a machine learning model Tome^42^). Environmental sources were also retrieved from the genome reference databases or from the associated published study (Table S1). As detailed current phylogeny of Thaumarchaeota is based on the *amoA* or 16S rRNA genes^2, 17, 43^, these two phylogenetically congruent markers were retrieved using RNAmmer^44^ or BLASTp^45^ against two representative *amoA* databases^2, 17^. Their classification towards the most up-to-date Thaumarchaeota *amoA* and 16S rRNA gene databases^2, 43^ was performed using BLASTn^45^. Initial classification of genome sequences was performed using classify_wf in the GTDB-Tk package^46^.

### Phylogenomic reconstruction and proteome content changes across evolutionary history

#### Datasets

Two datasets were used in this study: a phylum-level phylogenomic reconstruction including 165 genomes with a completeness greater than 45 % and contamination lower than 10%, and Nitrososphaerales-level phylogenomic reconstruction performed on the 26 genomes belonging to the lineage.

#### Ortholog selection

The best of three ortholog selection methodologies was selected (see Supplementary Information: Extended phylogenomics). Ortholog groups (OGs) were detected using Roary (-i 50, -iv 1.5)^47^ and core OGs were defined as those that were present in only one copy in each genome and were present in at least 85 % of the near complete genomes. Core OGs were aligned individually using MAFFT L-INS-i^48^ and spurious sequences and poorly aligned regions were removed with trimal (automated1, resoverlap 0.55 and seqoverlap 60)^49^. Alignments were removed from further analysis if they presented evidence of recombination using the PHItest^50^. Such ortholog prediction using the MCL algorithm resulted in 75 and 188 marker genes for the phylum-level and the Nitrososphaerales-level phylogenomic reconstruction, respectively (workflow illustrated in Figure S5).

#### Phylogenomic tree construction

Phylogenomic reconstruction was performed on each dataset, using a concatenated supermatrix of core OG alignments. Each maximum likelihood tree was constructed with IQ-TREE^51^ using the best fitting protein model predicted in ModelFinder^52^ and an edge-linked partition model (model details in Table S13). Branch supports were computed using the SH-aLRT test^53^ and 2,000 ultrafast bootstraps, further optimised with a hill-climbing nearest neighbor interchange (NNI) search.

#### Proteome content changes across evolutionary history

For the phylum-level dataset, protein families were detected with Roary with reduced stringency (-i 35, –iv 1.3, –s) and sequences shorter than 30 amino acids and families with less than 4 sequences were removed from further analysis. All remaining sequences within each family were aligned using MAFFT L-INS-I, processed with trimal (automated1) and ML phylogenetic trees were constructed for each alignment (IQ-TREE, -bb 1000, -bnni, -MFP). Ninety-seven percent (5,683 of the 5,850) of the protein family trees could be probabilistically reconciled against the supermatrix tree using the ALEml_undated algorithm of the ALE package^16^ to infer the numbers of duplications, losses, intra-LGT (transfer within the sampled genome set) and originations (including both inter-LGT, e.g. transfer from other phyla outside the sampled genome set or *de novo* gene formation) on each branch of the supermatrix tree. For the purposes of this analysis the small number of genes transferred between the related phyla Thaumarchaeota, Aigarchaeota and Bathyarchaeota genomes studied here are deemed intra-LGT events, not origination events. Ancestral proteome content (both size and predicted metabolic pathways) was estimated for each node of the tree. All phylogenomic trees were visualised using FigTree (http://tree.bio.ed.ac.uk/software/figtree/) and iTOL^54^.

### Taxonomic affiliation

Internally consistent taxonomic levels were calculated using an agglomerative clustering approach. Average amino acid identities (AAIs) between pairwise sets of genomes were calculated using CompareM (https://github.com/dparks1134/CompareM) and AAIs hierarchical clustering was performed in MeV (https://sourceforge.net/projects/mev-tm4/) using Euclidean distance with complete linkage. Maximum divergence is defined as the lowest AAI between any two members of the same group, subtracted from 100%. Pairwise relative phylogenetic distances between each genome were subjected to hierarchical clustering using single linkage Pearson correlation and taxonomic levels corresponding to class, order and family were defined as Pearson’s distances of less than 0.34, 0.13 and 0.015, respectively. These cut-off distances were chosen by comparison to existing thaumarchaeotal taxonomy^5-7, 55-63^. The taxonomic levels for genus and species levels were defined with AAI thresholds higher than 70% and 95%, respectively^64, 65^. The nomenclature of the different taxonomic stratifications relied primarily on previously existing classifications, with attribution of the names being primarily based on the strains whose cultivation was first reported. For clusters with no reported culture, the family-level nomenclature inferred from previously ammonia monooxygenase phylogenetic tree^2^ was used. For groups not matching those affiliations, such as genomes lacking the ammonia monooxygenase, the first listed genome of the group was used to name the group. In addition, the topology of a subset of the phylogenomic tree (focusing on ammonia-oxidising Thaumarchaeota) was compared to previously published *amoA*-based phylogenies^2, 13^ at the family level.

### Functional annotation and metabolic reconstruction

For each protein family, a medoid sequence (the sequence with the shortest summed genetic distances to all other sequences in the family) was calculated under the BLOSUM62 substitution matrix using DistanceCalculator in Phylo (https://biopython.org/wiki/Phylo). Medoids were annotated against the KEGG database^66^ using GhostKOALA^67^, against the arCOG database^68^ using Diamond BLASTp^69^ (best-hit and removing matches with e-value > 10^−5^, % ID < 35, alignment length < 80 or bit score < 100) and against the Pfam database^70^ using hmmsearch^71^ (-T 80). Carbohydrate active enzymes were annotated using HMM models from dbCAN (http://bcb.unl.edu/dbCAN2/) (filtered with hmmscan-parser.sh and by removing matches with mean posterior probability < 0.7). The predicted protein families present within each ancestor genome was inferred from the ALE results, allowing for ancestral metabolic reconstruction at each node of the phylogenetic tree. Proteome novelty in extant genomes was defined as the percentage of proteins encoded by a genome that do not have a homolog in the arCOG database. Origins of laterally acquired gene families were estimated by querying the medoid against UniRef90^72^ sequences with strain-level designations and excluding thaumarchaeotal matches.

## Acknowledgments

This work and P.O.S. were financially supported by UKRI through the NERC grant (NE/R001529/1). In addition, C.G.-R. and T.A.W. were both supported by a Royal Society University Research Fellowship (URF150571 and UF140626). C.Q. was funded through an MRC fellowship (MR/M50161X/1) as part of the CLoud Infrastructure for Microbial Genomics (CLIMB) consortium (MR/L015080/1). S. R. was funded through the BBSRC award ‘EBI Metagenomics - enabling the reconstruction of microbial populations’ BB/R015171/1. The Thames Metagenome Database was funded through the NERC award ‘Using next generation sequencing to reveal human impact on aquatic reservoirs of antibiotic resistant bacteria at the catchment scale’ (NE/M011674/1, NE/M011259/1, NE/M01133X/1).

We thank Tony Travis for his support with Biolinux and acknowledge Jim Prosser for his critical reading of the manuscript. The authors would also like to acknowledge the support of the Maxwell computer cluster funded by the University of Aberdeen.

## Author contributions

P.O.S., T.A.W. and C.G.-R. designed the study and developed the theory. C.Q. and S.R. assembled the 12 new metagenomes. The Thames consortium collected samples and performed DNA extraction for the metagenome assembly. P.O.S., T.A.W. and C.G.-R. interpreted the results and wrote the manuscript. All authors accepted the final version of the manuscript.

## Thames Consortium

William H Gaze^4^, Jennifer Holden^5^, Andrew Mead^6^, Sebastien Raguideau^3^, Christopher Quince^3^, Andrew C Singer^7^, Elizabeth M H Wellington^5^, Lihong Zhang^4^

^3^Warwick Medical School, University of Warwick, UK. ^4^European Centre for Environment and Human Health, Medical School, University of Exeter, UK. ^5^School of Life Sciences, University of Warwick, UK. ^6^Rothamsted Research, Harpenden, UK. ^7^UK Centre for Ecology & Hydrology, Wallingford, OX10 8BB, UK.

## Code availability

Custom scripts for manipulating ALE outputs have be deposited at https://github.com/Tancata/phylo/tree/master/ALE.

## Data availability

Genome sequences assembled in this work are available from Genbank under the accession numbers JAATVI000000000 (Nitrososphaerales archaeon TH1173), JAATVJ000000000 (Nitrososphaerales archaeon TH1177), JAATVK000000000 (Nitrososphaerales archaeon TH5894), JAATVL000000000 (Nitrososphaerales archaeon TH703), JAATVM000000000 (Nitrososphaerales archaeon TH1920), JAATVN000000000 (Nitrososphaerales archaeon TH5888), JAATVO000000000 (Nitrososphaerales archaeon TH4187), JAATVP000000000 (Nitrososphaerales archaeon TH5896), JAATVQ000000000 (Nitrososphaerales archaeon TH1917), JAATVR000000000 (Nitrososphaerales archaeon TH526), JAATVS000000000 (Nitrososphaerales archaeon TH5895) and JAATVT000000000 (Nitrososphaerales archaeon TH5893).

## Notes

https://figshare.com/s/fc84c2a9f1052c833858

### Competing Interest Statement

The authors have declared no competing interest.

