## Supplementary material for "Gene duplication drives genome expansion in Thaumarchaeota"

### Material and methods

#### *Sampling, sequencing and assembly methodology*

Samples were taken from 23 sites spread across 7 tributaries of the Thames river at three different times, with duplicate samples taken in the summer of 2015 and triplicate samples taken in the summer and winter of 2016. DNA extraction was performed from 0.5 g of sample within 12 h of collection using the FastDNA™ spin kit for soil, following manufacturer's protocol. Libraries were constructed with TruSeq® DNA library preparation kit and sequenced on the HiSeq2500 platform (2 x 150 bp reads). Sequences were obtained from 171 samples at an average read depth of 45 million, resulting in 2.3 Tbp. The 171 samples were co-assembled using MEGAHIT[1] with default parameters giving a 59.3 Gbp assembly with an N50 of 999 bp and a total of 4,340,028 contigs greater than 2,000 bp in length. Reads from every sample were mapped back onto these contigs using bwa-mem[2] with default parameters and coverage depths were calculated. These coverage depths and the sequence composition of the contigs were used by the CONCOCT clustering algorithm[3] to produce 6,008 genomic bins. Twelve of theses bins were found to possess at least one of the *amo*A or *amo*B genes that are characteristic of ammonia oxidizing archaea, detected using BLASTn[4] against a custom database of *amo*A and *amo*B sequences[5].

#### *Extended Phylogenomics*

*Estimating phylogeny:* In order to establish a robust phylogeny of the Thaumarchaeota, three methods of phylogenetic marker gene selection were used. In the first method, 43 marker genes were selected using CheckM[6]. One ortholog group was removed, as it presented evidence of recombination using the PHItest[7] and the remaining 42 genes were used as marker genes. In the second method, the Hidden Markov Model (HMM) profiles of the 122 archaeal phylogenetic markers described by GTDB[8] were downloaded from Pfam[9] and TIGRFAMS[10] and searched for in the genomes using hmmsearch[11] (-T 80). For the 80 matching proteins that were present in single copy in at least 70% of the genomes, the regions aligning to the HMM profiles were extracted and used as marker genes. In the third method, orthologs were detected using the MCL algorithm-based software, Roary[12] (-i 50, -iv 1.5) and core ortholog groups were defined as those that were present in only one copy in each genome and were present in at least 85 % of the genomes presenting more than 80 % completeness and less than 5 % contamination (workflow illustrated in Figure S5). No evidence of recombination in these orthologs groups was detected using the PHItest, resulting in 75 marker genes.

For all three datasets, marker genes were aligned individually using MAFFT L-INS-i[13] and spurious sequences and poorly aligned regions were removed with trimal[14] (automated1, resoverlap 0.55 and seqoverlap 60). A maximum likelihood tree was constructed for a concatenated supermatrix of alignments with IQ-TREE[15] using the best fitting protein model in ModelFinder[16] for each alignment and an edge-linked partition model. Branch validation involved SH-aLRT test[17] and 2000 ultrafast bootstraps, further optimised with a hill-climbing nearest neighbor interchange (NNI) search.

*Comparing phylogeny estimations:* Likelihoods for each of the phylogenies were calculated by amalgamated likelihood estimation of 5,683 thaumarcheotal gene families and used to perform an approximate unbiased tested of the three trees in the manner described by Williams *et al*

2017[18] and implemented in ALE[19] and CONSEL[20]. The most likely tree was used for further analysis.

<u>*Topology testing in constraint trees:*</u> Topology tests were performed with IQ-TREE using the MCL markers approach. The aim was to compare an unconstrained MCL tree with three constrained trees that represent the major incongruences between this work and the phylogenomic tree proposed by Ren *et al.* [21]: a) the constrained tree 1 (Monophyletic yellow, using the colour scheme from Figure S6) forces FN1, YP1-bin3 and UBA183 to form a monophyletic clade with AD-613-B23, RBG-16-49-6, SAT139, EAC691, UBA57, UBA223 and SP3992; b) the constrained tree 2 (Paraphyletic blue) forces 3F and SCU2 to form a paraphyletic clade with SAT137 and UBA213; c) the constrained tree 3 (Monophyletic yellow and paraphyletic blue) possesses the constraints of both constrained tree 1 and 2.

### Results

#### *A robust phylogeny for the Thaumarchaeota*

A phylogenomic approach was adopted to explore the relationships of 152 Thaumarchaeota genomes, 11 Aigarchaeota and 2 Bathyarchaeota genomes. As the use of different phylogenetic markers and tree construction methods has been known to produce dramatically different estimates of phylogeny, we examined three different phylogenomic marker sets and compared the inferred topologies.

The tree constructed using CheckM markers is highly similar to the phylogeny previously proposed by Ren *et al* [21]. The HMM and MCL are largely congruent with these trees, albeit with some differences. In the CheckM tree, SAT137 and UBA213 form a separate clade to J079, 3F and SCU2, whereas these organisms for a single clade in the HMM and MCL trees (Figure S6, blue dots). In the HMM and MCL trees, FN1, YP1-bin3 and UBA183 form a separate clade to AD-613-B23, RBG-16-49-6, SAT139, EAC691, UBA57, UBA223 and

SP3992, whereas using the CheckM markers these organisms form a single clade (Figure S6, yellow dots). In the MCL tree, BS3, BS4, ARK01, UBA164, DS1 and UBA160 form a strongly supported monophyletic group. This group is also present in the HMM tree (albeit with poor support) and is split into two poorly supported groups in the CheckM tree (Figure S6, red dots). The three trees gave incongruent topologies for the Nitrososphaerales (Figure S6, green dots), so we performed a targeted analysis of this order using more phylogenetic markers, as discussed in the main text. The resulting topology was consistent with that obtained using the MCL markers in the phylum-wide analysis.

As the three trees are providing strong support (SH-alrt and ultra-fast bootstrap) for some contradicting phylogenies, an additional tree verification test was performed. Different species tree estimations imply different scenarios of gene family evolution, resulting in different gene family likelihoods when using a probabilistic gene tree to species tree reconciliation model. It was therefore possible to use an approximately unbiased test to establish a confidence values for the tree alternative phylogeny estimations. From this analysis, it was possible to reject all but the MCL tree (Table S14).

Additionally, topology tests were performed using the MCL markers that compared the unconstrained MCL tree to three constrained trees that represented two of the incongruent placements observed between the CheckM tree and the other two trees, namely splitting of the blue dotted clade and merging of the yellow dotted clades from Figure S6. The combination of monophyletic yellow clade and paraphyletic blue clade could be significantly rejected by approximate unbiased testing and several other tests (Table S15). While the individual incongruences could not be statistically rejected by this method, they were far less favoured than the unconstrained MCL tree topology by all statistical tests applied here (Table S15).

Finally, as part of the phylogenomic workflow described in Figure S5, a species tree was formed using the same marker genes but including only the 87 genomes sequences of greater than 80 % completeness and less than 5 % contamination. The resulting tree is largely in agreement with the full 165 genome species tree, with no strongly supported contradictions (Figure S7). This indicates that missing data in the 165 genome supermatrix, resulting from some genomes lacking some of the markers, has not had a major effect on the topology of the species tree.

***Extended optimal growth temperature***

*In silico* prediction of optimal growth temperature was performed with Tome, a machine-learning model that uses 2-mer amino acid composition across an organism proteome. Results obtained from this analysis were validated against a set of Thaumarchaeota for which the optimal growth temperature has been experimentally determined. Tome proved to be relatively accurate in predicting optimal growth temperature for the mesophilic (20-40 °C) Thaumarchaeota, with an average difference of 3 °C between the experimental and *in silico* estimations (Table S16). However, this difference increases to an average of 10 °C in the thermophiles (41-122 °C) (Table S16), even if predicted values were related to experimental values ($R^2$=0.8744). Therefore, Tome can reasonably predict the mesophilic or thermophilic state of an organism from the genome, but optimal growth temperatures may be significantly higher than predicted in thermophiles.

**Supplementary Figures.**

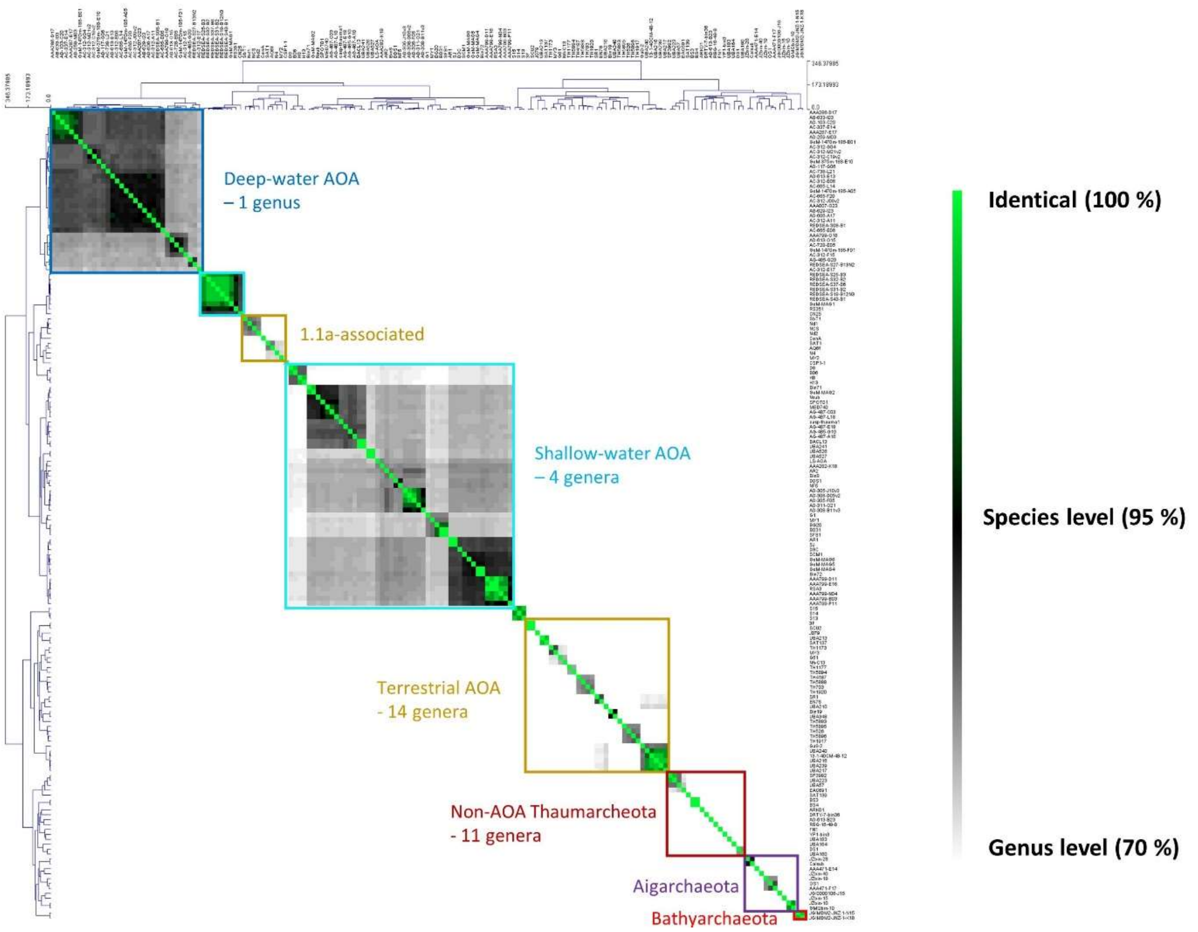


**Figure S1. Average amino acid identity of Thaumarchaeota and related archaea.** Average amino acid identity (AAI) was used as a measure of genome similarity. Pairwise AAIs were clustered using Euclidean distance with complete linkage. AAIs were coloured at the species level (> 95 %) from green to black and at the genus level (> 70 %) from black to light grey. AAI values below 70% are coloured white.

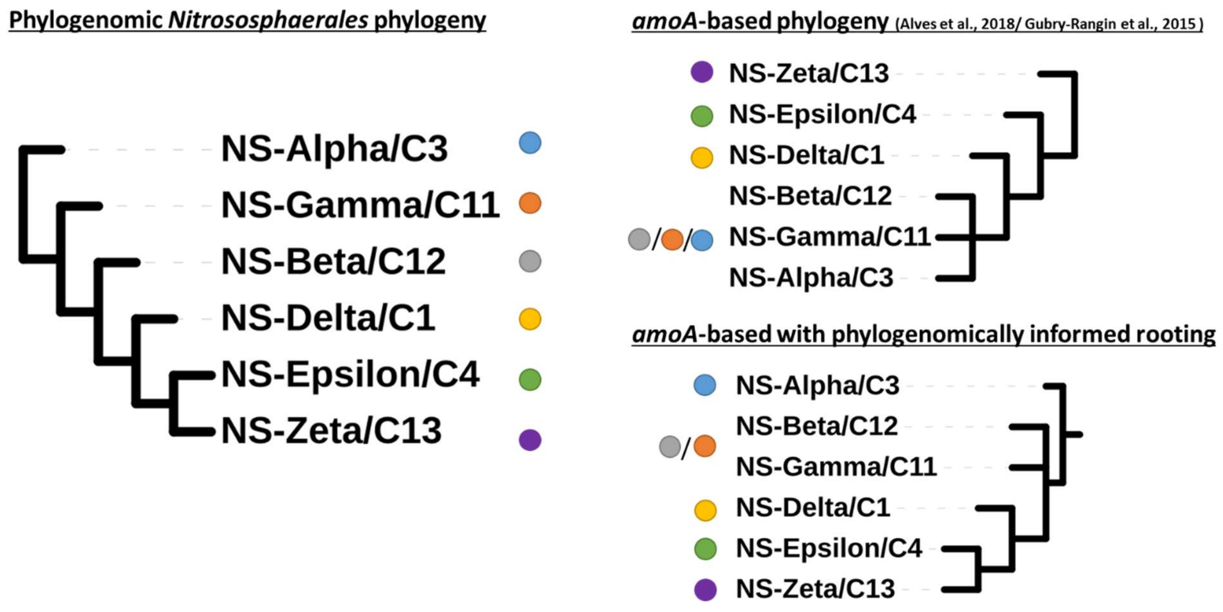


**Figure S2. Phylogenomic vs *amoA*-based phylogeny of Nitrososphaerales.** For the phylogenomic and the Alves *amoA*-based phylogenies[22], branches with less than 95% ufboot and 85% SH-aLRT were collapsed, while branches with less than 75% posterior probability were collapsed for the Gubry-Rangin *amoA*-based phylogeny[23]. Both *amoA*-based phylogenies were identical at family level. Coloured circles are used to emphasise the family-level groups. Slashes (/) indicate were branches could not be confidently resolved.

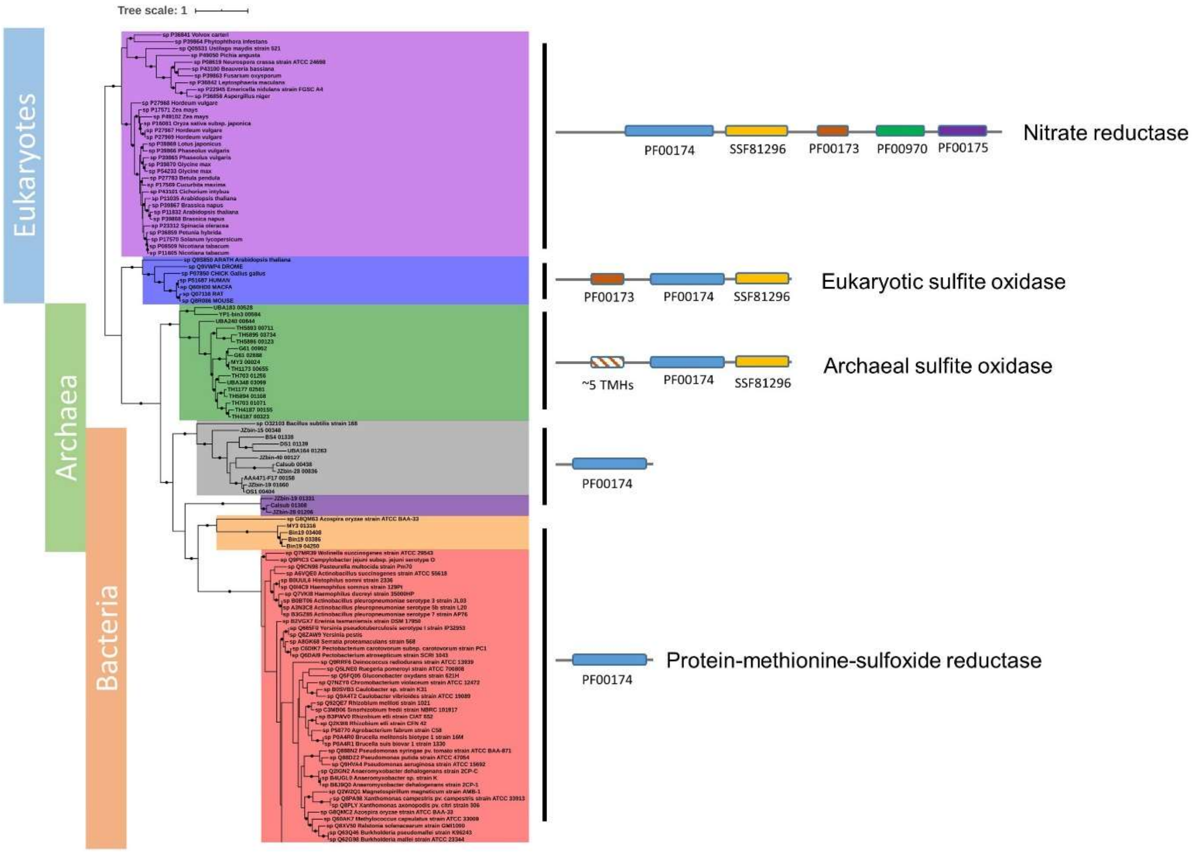


**Figure S3. Sulfite oxidase family molybdoenzymes of Thaumarchaeota.** MoZymes of Thaumarchaeota represent a putative archaeal class of sulphite oxidases. Dots indicate branches with greater than 95% of 2,000 ultrafast bootstraps. Swiss-Prot sequences possessing the PF00174 domain were used as references. Coloured bars on left indicate the domain of origin of protein clades. Overlapping of these bars indicates clades with proteins from more than one domain of life. Transmembrane helices (TMHs). Figures to the right of tree are schematic representations of domain organisation in corresponding protein clade.

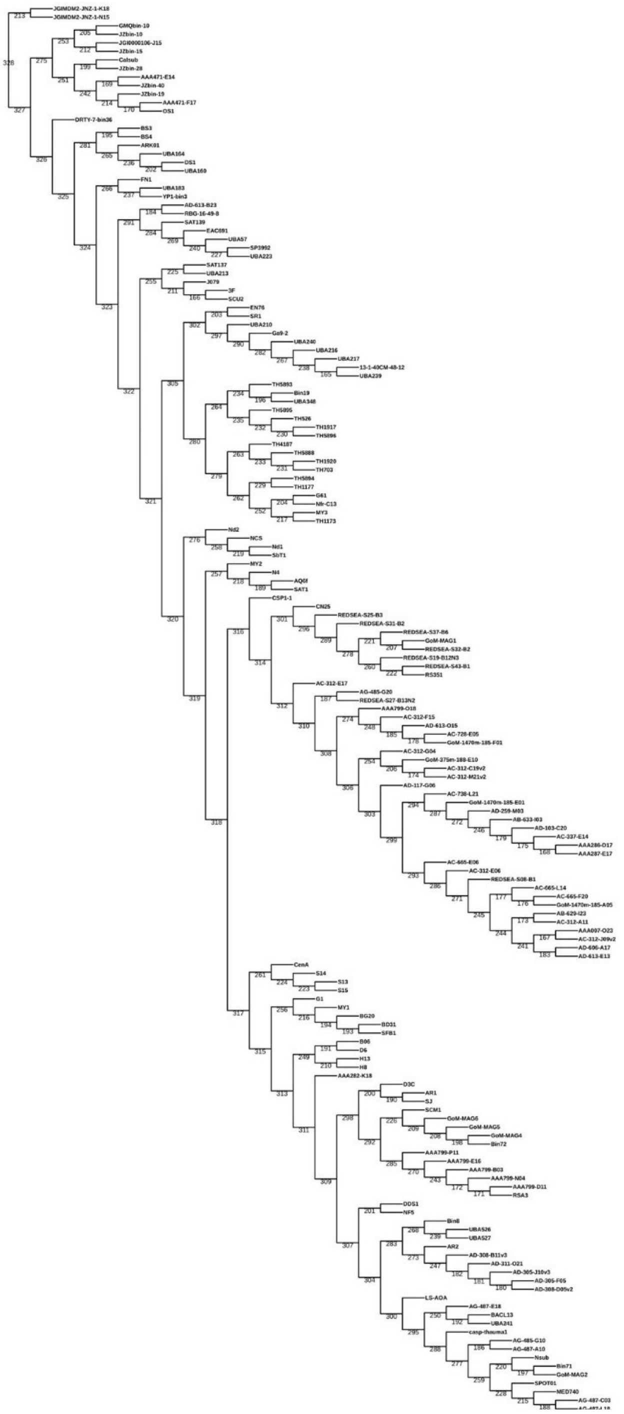

**Figure S4. Branch labelled phylogeny of Thaumarchaeota and related species.** This cladogram possesses the topology of the tree presented in Figure 1. The tree branch numbers corresponded to the quantification of gene content change mechanisms in Table S4. These two datasets were combined to form Figure 3.

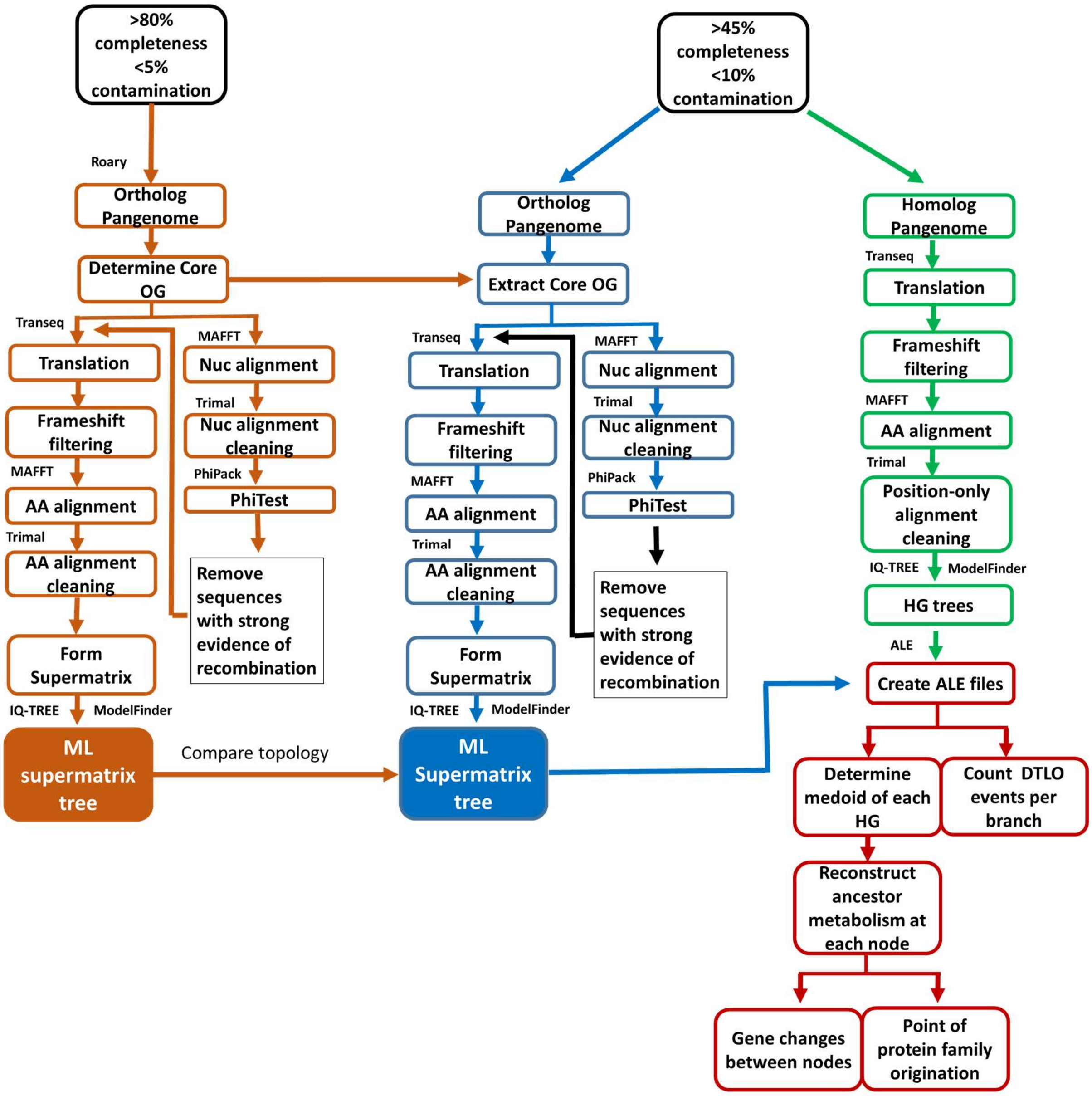


**Figure S5. Schematic workflow of phylogenomic and evolutionary analysis.** This workflow describes the steps taken in the construction of the ML tree presented in Figure 1 and in gene family history prediction used to construct the Figure 2. Tools used in the orange, blue and green branches of the workflow have been shown next to their stage of use. Custom scripts for manipulating ALE outputs have be deposited at https://github.com/Tancata/phylo/tree/master/ALE .

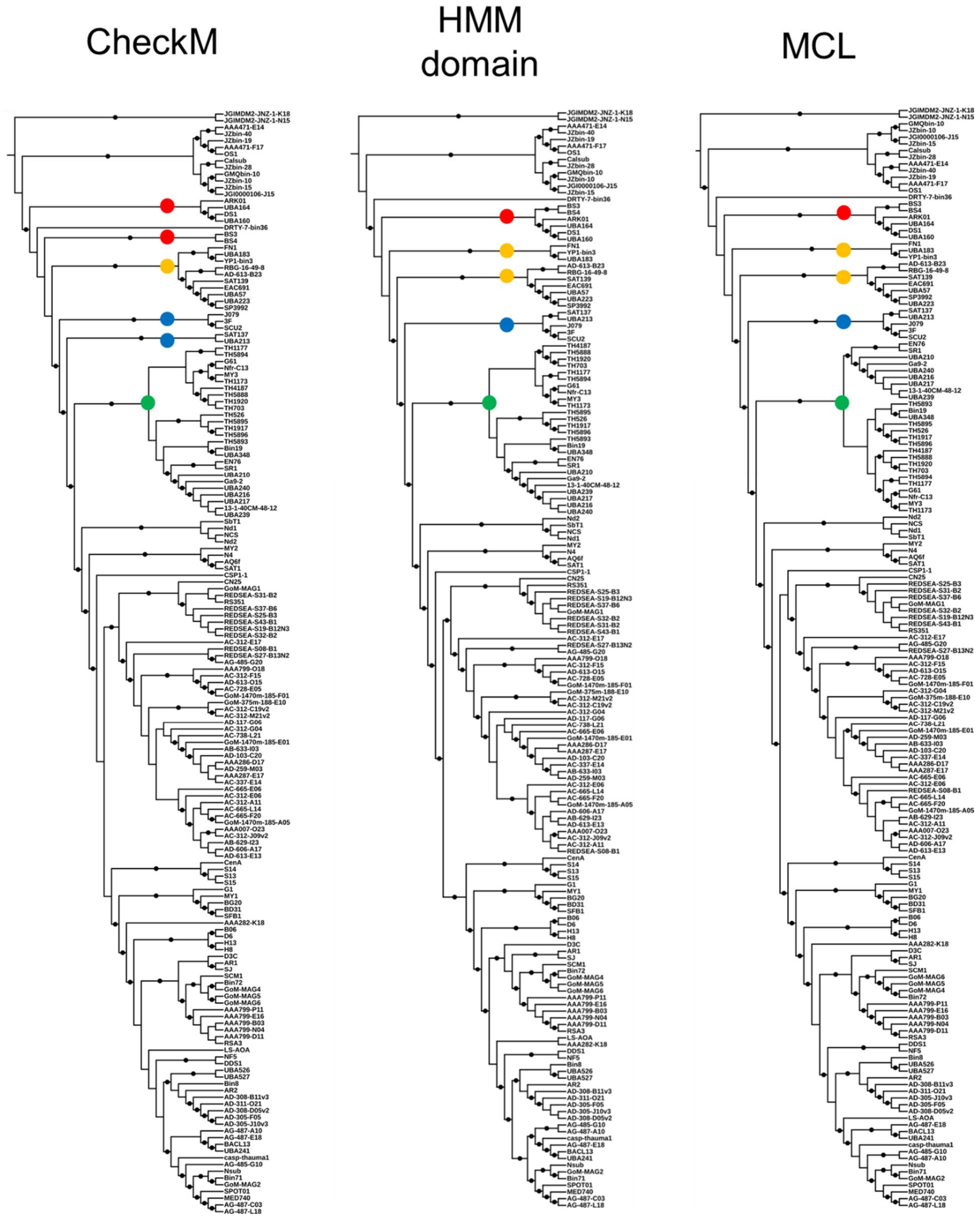


**Figure S6. ML trees using 3 methods of phylomarker selection.** Coloured circles are used to highlight areas of incongruence between methods. Dots indicate branches with greater than 95% of 2,000 ultrafast bootstraps and 1,000 SH-alrts. The tree is rooted with the Bathyarchaeota strains, JGIMDM2-JNZ-1-K18 and JGIMDM2-JNZ-1-N15.

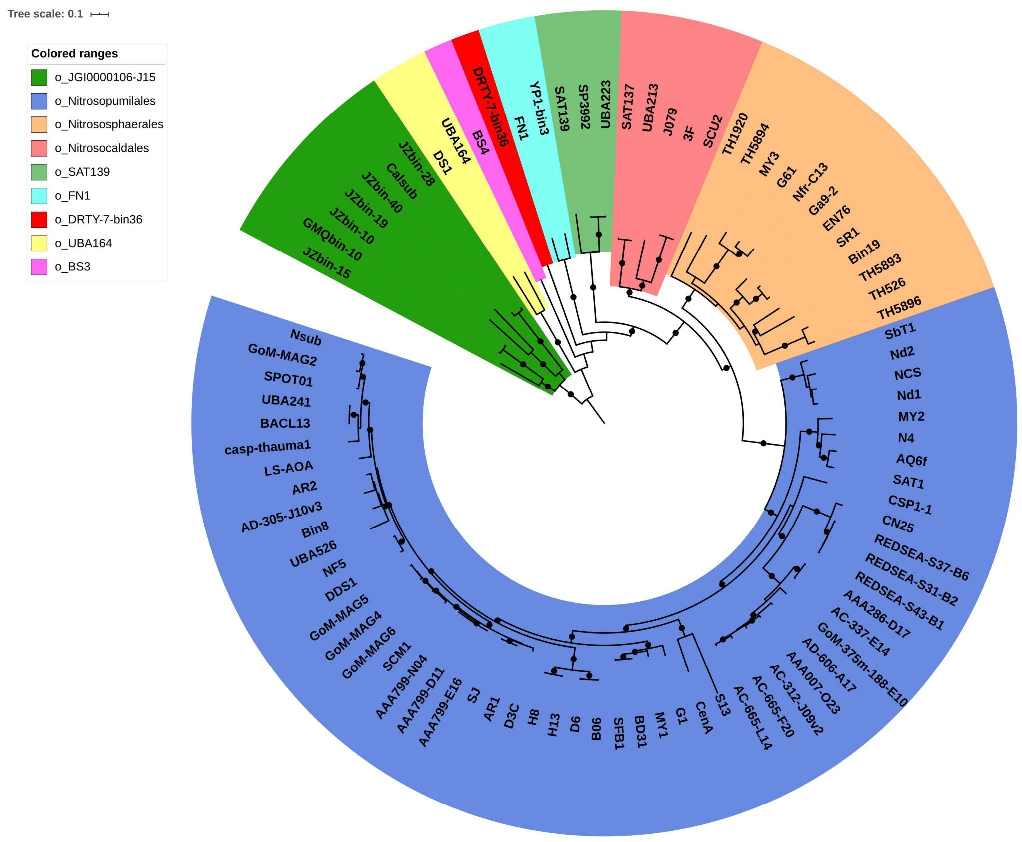


**Figure S7. Phylogeny of near complete Thaumarchaeotal and related archaeal genomes.** ML tree of 75 concatenated ortholog groups. Dots indicate branches with greater than 95% of 2,000 ultrafast bootstraps and 1,000 SH-alrts. Colours indicate order-level groups. The tree is rooted with the Aigarchaeota strains (o_JGI0000106-J15).

**Legends of supplementary tables**

**Table S1. Genome descriptions.** Metagenome assembled genome (MAG), single-cell assembled genome (SAG). Protein novelty is defined as the percentage of encoded proteins that lack a close homolog (e-value < 10-5, % ID > 35, alignment length > 80 and bit score > 100) in the arCOG database.

**Table S2. Taxonomic stratification of Thaumarchaeota.** Grey columns display the closest match of query *amo*A and 16S rRNA genes to sequences in Thaumarchaeota phylogenetic databases[5, 22].

**Table S3. GTDBTk classify_wf classification.** Relative evolutionary divergence was calculated using the Genome taxonomy database toolkit available at https://github.com/Ecogenomics/GTDBTk.

**Table S4. Quantified mechanisms of proteome change.** Gene content changes on each branch of the phylogenomic tree presented in Figure 1 have been divided into four evolutionary mechanisms of change: duplications, losses, intra-LGT and originations. Duplication and loss are defined as the copying and loss of a gene within a genome, respectively. Intra-LGT is defined as the acquisition of a gene from other member(s) of the phylum, while originations are defined as the acquisition of a gene from members of other phyla outside the sampled genome set and possibly by de novo gene formation.

**Table S5. arCOG copy number gain at duplication hotspots.** The increase in copy number of particular arCOG families between the query genome reconstruction and its last ancestor.

**Table S6. arCOG copy number loss at duplication hotspots.** The decrease in copy number of particular arCOG families between the query genome reconstruction and its last ancestor.

**Table S7. arCOG annotations of originating gene families.** Letters in second column represent clusters of orthologous groups (COG) categories.

**Table S8. Best hit taxonomy of originating gene families.** Best results obtain by querying the protein family mediod against UniRef90 sequences with strain-level designations and excluding thaumarchaeotal matches.

**Table S9. Duplication and losses in originating and ancestral gene families.** The percentage of gene families that have duplicated or been lost in gene families that have originated on LNS (Branch 305), LNS-2 (Branch 280) and prior to LNS (ancestral gene families).

**Table S10. Key pathways in Thaumarchaeota genomes and ancestral reconstructions.** Describes the presence (black) or absence (white) of selected genes in ancestral reconstructions. Numbers in columns of extant genomes represent the copy number of the corresponding gene.

**Table S11. Functional gains in major evolutionary transitions.** KEGG k numbers present in the stated descendant and absent in the stated ancestor.

**Table S12. Functional losses in major evolutionary transitions.** KEGG k numbers absent in the stated descendant and present in the stated ancestor.

**Table S13. Best fitting model of core orthologs groups.** The best fitting sequence evolution model for each partitioned gene used in the phylogenomic tree estimation in Figure 1. This was determined using ModelFinder[16].

**Table S14. Comparing different species tree estimations using gene tree-species tree reconciliation.** The most likely species tree was predicted using the following tests: Log-likelihood difference (logL), approximate unbiased test (AU), bootstrap probability calculated from the multiscale bootstrap (NP), bootstrap probability calculated in the usual manner (BP),

Kishino-Hasegawa test (KH), Shimodaira-Hasegawa test (SH), weighted Kishino-Hasegawa test (WKH) and weighted Shimodaira-Hasegawa test (SH).

**Table S15. Comparing different species tree estimations using constrained trees.** The most likely species tree was predicted using the following tests: Log-likelihood difference (logL), logL difference from the maximal logl in the set (deltaL), bootstrap proportion using RELL method (RELL), Kishino-Hasegawa test (KH), Shimodaira-Hasegawa test (SH), weighted Kishino-Hasegawa test (WKH), weighted Shimodaira-Hasegawa test (WSH), expected likelihood weight (ELW), approximate unbiased test (AU) and can a tree be statistically rejected? (Rejected).

**Table S16. Comparison of genome-predicted and experimentally determined optimal growth temperatures in Thaumarchaeota.** Experimentally determined optimal growth temperatures were extracted from published literature[24-38] and *in silico* predicted optimal growth temperature was calculated using Tome[39].
